# Pseudorabies virus causes splenic injury via inducing the oxidative stress and apoptpsos related factors in mice

**DOI:** 10.1101/2023.09.01.555967

**Authors:** Wei Sun, Shanshan Liu, Yi Yan, Qingyan Wang, Yu Fan, Samuel Kumi Okyere

## Abstract

Pseudorabies virus (PRV) is an immunosuppressive disease that causes significant damage to the pig industry. This study aimed to detect the effects of PRV on oxidative stress related factors and cell apoptosis in the spleen, providing a basis for the research on the pathogenesis of PRV in mice model. Pathological observation was performed by hematoxylin and eosin Y staining. Biochemical and Flow cytometry method were performed to determine the reactive oxygen species profile of the spleen post-infection and apoptosis detection. In addition, q-PCR and Western blot were adopted to measure the apoptotic conditions of the spleen infected with PRV. The results indicated that the ROS level in the PRV infection group was remarkedly increased (*p*<0 01) at a time-dependent pattern. Furthermore, the Malondialdehyde levels in the spleen of mice in the infection group increased significantly (*p*<0.01) in a time-dependent mode. However, the Catalase, Superoxide dismutase, and Glutathione activity and expression levels in the infection group were significantly decreased with the control group (*p*<0 01) in a time-dependent manner. Furthermore, the ratio of splenocyte apoptosis in the infection group significantly increased (*p*<0 05, *p*<0 01) in a time-dependent manner. In conclusion, PRV infection causes apoptosis of the spleen via oxidative stress in mice.

## 1. Introduction

Pseudorabies virus (PRV), a dsDNA genome, belongs to the α members of the herpesvirus subfamily and is known to the pathogen of pseudorabies (PR)[1]. PR is a contagious disease that shows neurological symptoms and reproductive disorders. PR causes pathological changes such as inflammation and necrosis in parenchymal organs such as the brain, liver, spleen, and kidneys, and the degree of damage to these organs varies among different strains^2^. Studies have demonstrated that PRV disrupts the balance of cellular oxidation and antioxidant, along with the imbalance between reactive oxygen species (ROS) and their intermediates, inducing oxidative stress in cells, causing to the steatolysis, proteolysis, nucleic acid degradation in cells, triggering to cell apoptosis^9^, inflammatory response^10^ and immune suppression^11^. PRV infects various mammals, including ruminants, herbivores, and rodents hence, which brings huge losses in economy to the global pig industry^7^. However, pigs are the only natural reservoir of PRV[2]. Studies have shown that humans may also be potential hosts of PRV at the genetic level, indicating that PRV can break through species barriers and cause infection in humans[3]. In addition, there have been several reports about encephalitis patient infected by PRV[4-6]. However, there is currently no effective method for treating PRV infection available.

During viral infection, the relative expression of virus-related genes and the activation of innate antiviral response systems leads to an increase in ROS and toxic byproducts of energy metabolism. The imbalance between ROS and antioxidants including superoxide dismutase (SOD), catalase (CAT), glutathione (GSH), and glutathione peroxidase (GSH-Px) in the body leads to oxidative stress, cell death, and tissue/organ damage. ROS and the resulting changes in cellular redox status become one of the inducing factors for cell apoptosis.

The spleen as an immune organ is one of the major targets of PRV, however, the molecular mechanisms involved in PRV infection and toxicity of the spleen are not well established in the literature. Therefore, in this work, the effects and molecular mechanism of PRV on spleen toxicity in mice was investigated, aiming to give a theoretical basis for the key pathogenesis study of PRV as well as reveal other therapeutic targets for PRV infections.

## 2. Materials and Methods

### 2.1 Reagents and animals

Six-week-old SPF BALB/c mice were selected for the experiment, that was bought from Da-shuo Experimental Animal Co., Ltd, China. The PRV-HLJ strain (MK080279.1) was provided by the Animal Pathogen and Pathological Morphology Innovation Team of the Harbin Institute of Veterinary Medicine, CAAS, and is preserved in our laboratory. ROS detection kits (88-5930-74) were bought from Thermo Fisher Scientific-CN; The apoptosis kit with Annexin V-FITC/PI double staining was bought from BD Company in the United States; 2×SYBR Green PCR Mastermix (RR820A) and reverse transcription kit (RR047A) were purchased from TAKARA company; Tissue RNA extraction kit (B518621) was purchased from Shanghai Sangong Biological Co., Ltd; T-SOD (A001-1-2), MDA (A003-1), GSH (A006-2), GSH-Px (A005), and CAT (A007-1) detection kits are all purchased from Nanjing Jian-cheng Biotechnology Research Institute.

### 2.2 Experimental design

Mice were fed in a room with 12 h light and dark cycle, and water and diet were taken freely. The indoor temperature was maintained between 22-24 ℃ and the humidity between 40-60%. 36 mice were randomly grouped into 2 groups, the control group (18) which was subcutaneously administered with 200 μL of PBS and the infection group (18) which received subcutaneously inoculated with a 200 μL dose of 1×10^3^ TCID50/100 μL PRV. 6 mice in each group were dissected at 48, 72, and 96 h post-infection (hpi), and their spleen tissues were taken after after ether anesthesia for further experiments.

### 2.3 Pathological observation

The fresh spleen in 2.2 was fixed in 4% formaldehyde for more than 24 h. Then, dehydrate and transparent the tissue blocks embedded in paraffin and cut them into 5 pieces μM thick. Typically, 4-5 µm slices were cut and fixed on a glass slide. Parafn was removed by xylene and alcohol. Observe and photograph the tissue structure of the tissue stained with hematoxylin and eosin Y (H.E) using a digital camera.

### 2.4 Detection of ROS by flow cytometry in the Spleen

Take 300 μL spleen homogenate was mixed with 1 μL DCFH-DA dye solution, and was incubated in the darkroom at 37 ℃ for 20 min. Next, the treated tissue above was centrifuged at 4 ℃ for 5 min, and then added with 1 mL PBS solution. The supernatant was discarded and resuspended into 400 μL PBS. ROS was detected by using flow cytometry.

### 2.5 Detection of ROS-related factors in the Spleen

The spleens were collected from each group and ground to form 10% tissue homogenate by using a mechanical method. Then, the homogenate of all tissue was centrifuged at 2500 rpm for 10 min on the ice. The corresponding reagent kits were used to detect the changes in cellular oxidative stress-related factors including MDA, SOD, GSH, GSH-Px, and CAT.

### 2.6 Detection of splenic cell apoptosis

Fresh spleen tissues were taken from each experimental group and cut into 1 mm^3^ size. The tissue homogenates were prepared mechanically by grinding and passing through a 300 mesh filter. The cell concentration was adjusted to 1×10^6^/mL following by centrifugation and resuspension. Apoptosis with Annexin-V-FITC double stained kit was applied to measure the influence of PRV on the apoptosis of mouse splenocytes.

### 2.7 RNA extraction and qRT-PCR analysis

Following by the manufacturer’s instruction,”Animal Total RNA Separation Kit” was used to separate total RNA from the spleen. RNA integrity was determined by 1.5% agarose gel electrophoresis. A PrimeScript RT kit was used to convert total RNA to cDNA for qRT-PCR analysis. Next, cDNA was amplified by a PrimScript RT reagent Kit on the LightCycle 96 device (Roche, Germany) following the protocol provided. All oligonucleotide primers for this study were designed by Primer software and synthesized at Sagon Biotech Ltd, China. All qRT-PCR reactions system were mixed with the SYBR staining. Detailed information on primers can be found in Table 1. All relative expression data was calculated by the 2^-ΔΔCt^ method. And β-actin was introduced as an internal reference gene in this study.

**Table 1.**
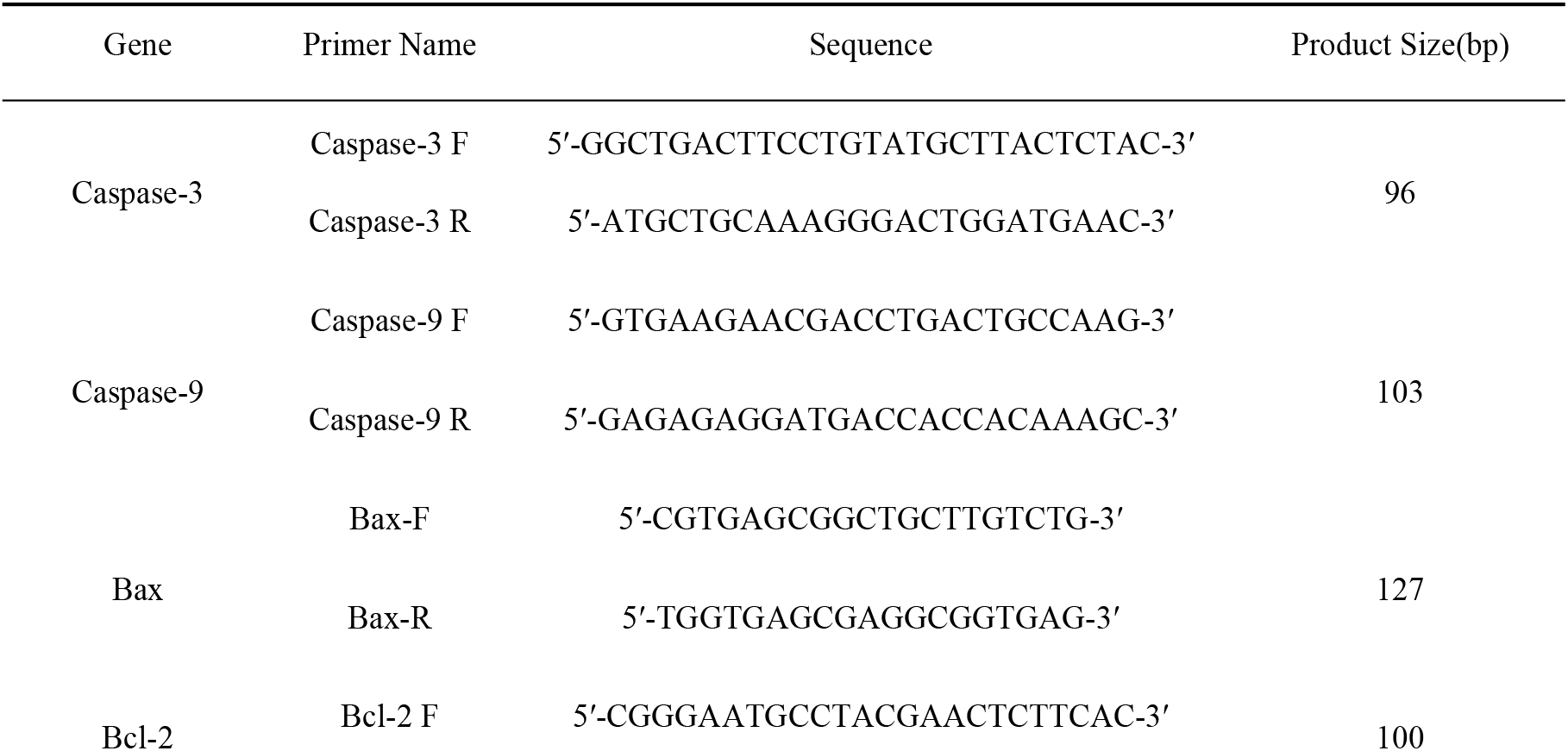
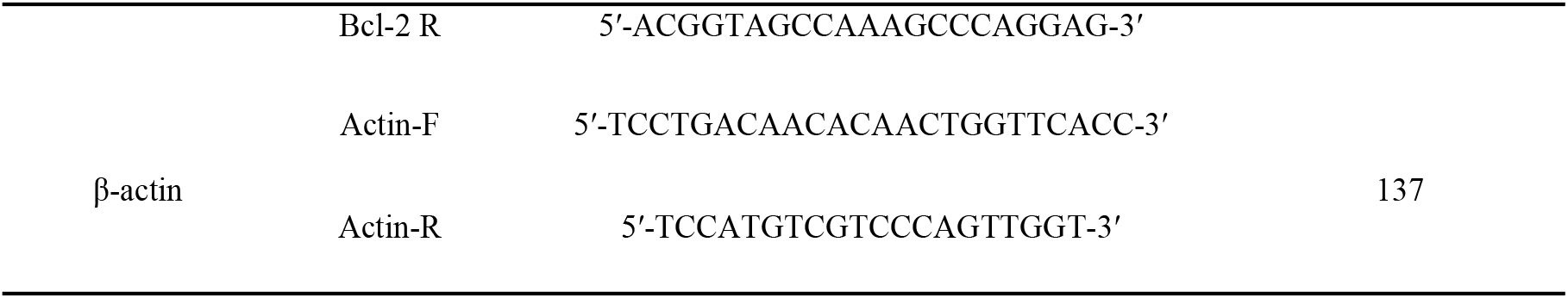
Primers information for qRT-PCR.

### 2.8 Western blot analyses

“Tissue Total Protein Extraction Kit” was used to extract of the total protein from spleen in all groups. Then, protein concentrations in each group were measured according to the Bradford method. Protein samples were first separated on 10% SDS-PAGE solution and the transferred onto PVDF membranes. Next, PVDF membranes wer incubated with the corresponding primary antibody at 4 ℃ for 12 h, i.e Caspase-3 (1:900), Caspase-9(1:1000), Bax (1:800), Bcl-2 (1:600) and β-actin (1:2000), followed by blocking in 5% skimmed milk solution for 2 h, respectively. The protein blot bands were washed three times with 5% PBST solution and reacted with HRP-labeled Goat anti-rabbit antibody for 1h with slight shaking. The membrane was observed on ECL after washing with PBST. The expression level of the proteins of interest to β-actin was calculated by using Using ImageJ2x software.

### 2.9 Statistical analysis

SPSS22.0 software was used to analyse the results, and the experimental data were expressed in the form of mean±standard deviation (mean±SD). One-way analysis of variance (ANOVA) algorithm was applied to evaluate the statistical differences among PRV infection and the control groups. The data histograms were plotted by GraphPad Prism 6.0 software. *p*<0.05 and *p*<0.01 value was introduced to determine the significant of differences.

## 3. Results

### 3.1 Pathological observation on Spleen

Based on the microscopic examination (Figure 1-A and B), the structure of the spleen tissue is normal, and no obvious histopathological damage is observed in the control and 48 hpi group. However, we observed different pathological changes in the red pulp area, mainly including congestion(Figure 1-C and D, yellow arrow) and even focal infiltration of neutrophils(Figure 1-D, green arrow), while no abnormalities were observed in the white pulp area.

**Figure 1.**
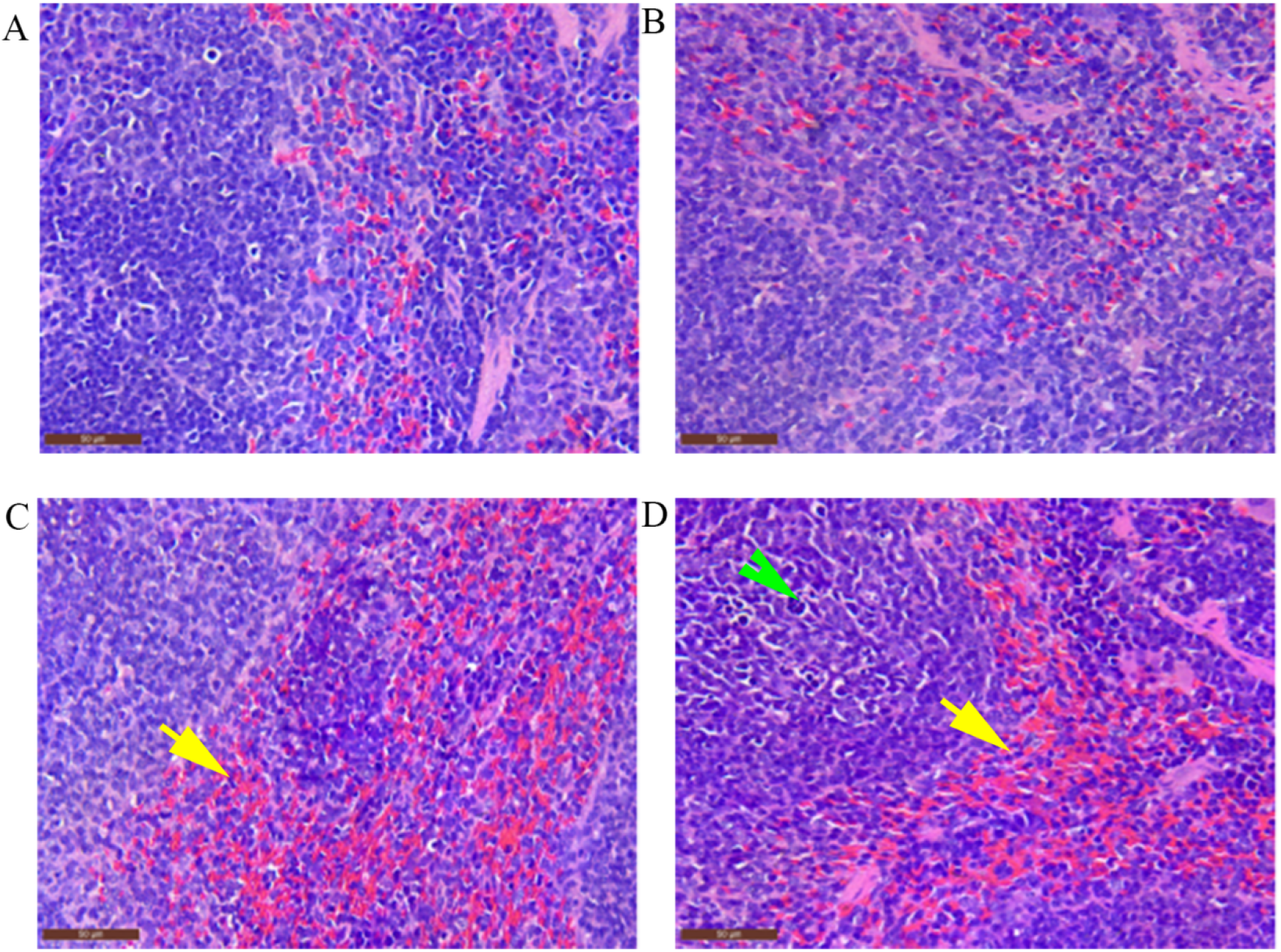
The pathological observation on Spleen(400×, Bar=50 μm) (A) Control; (B) 48 hpi; (C) 72 hpi; (D) 96 hpi

### 3.2 The influence of PRV on ROS levels in the Spleen

The results from the flow cytometry indicated that, compared with that in their respective control groups, ROS production in the spleen of the PRV infection group significantly increased in the whole infection period (*p*<0.01, Figure 2).

**Figure 2.**
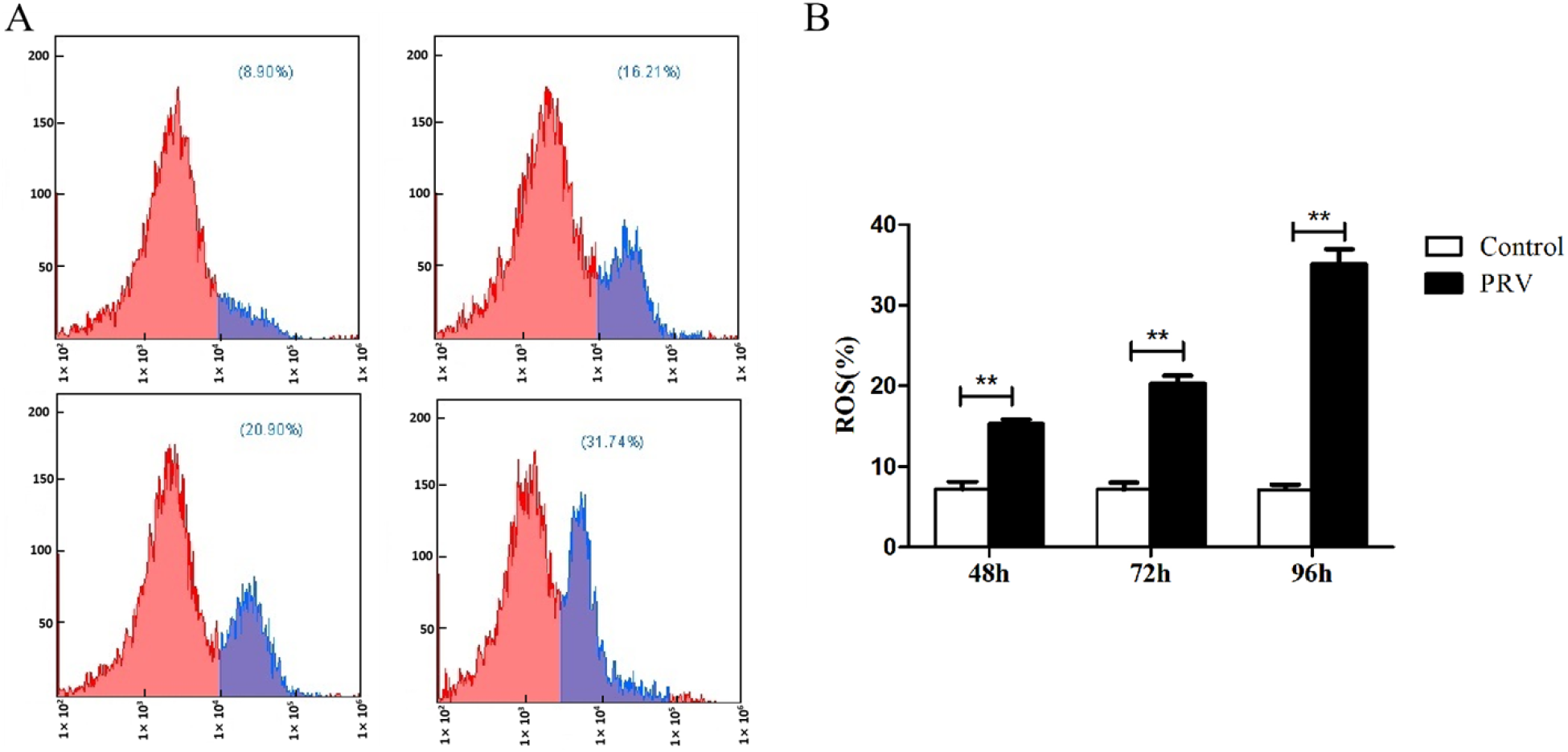
The inducement of PRV on production of ROS in the spleen. (A) The results of ROS detected by flow cytometry; (B) ROS statistical analysis histogram \*\**p*< 0.01 versus mock infection, the same as below.

### 3.3 The effect of PRV on oxidative stress-related factors in the spleen

As shown in Figure 3, the PRV treatment group significantly promoted the production of MDA in the spleen more than their control groups (*p*<0 01) in time dependent manner. However, PRV significantly descends the levels of CAT and GSH in the spleens in a time-dependant manner (*p*<0 05, *p*<0 01). In addition, compared with the control group, SOD and GSH-Px activities at the time point of 48 hpi showed no statistical difference (*p>*0.05). However, there was a significant decrease in both the levels of SOD and GSH-Px in the PRV infection group at the 72 hpi and 96 hpi time points compared to their respective controls (*p*<0 01).

**Figure 3.**
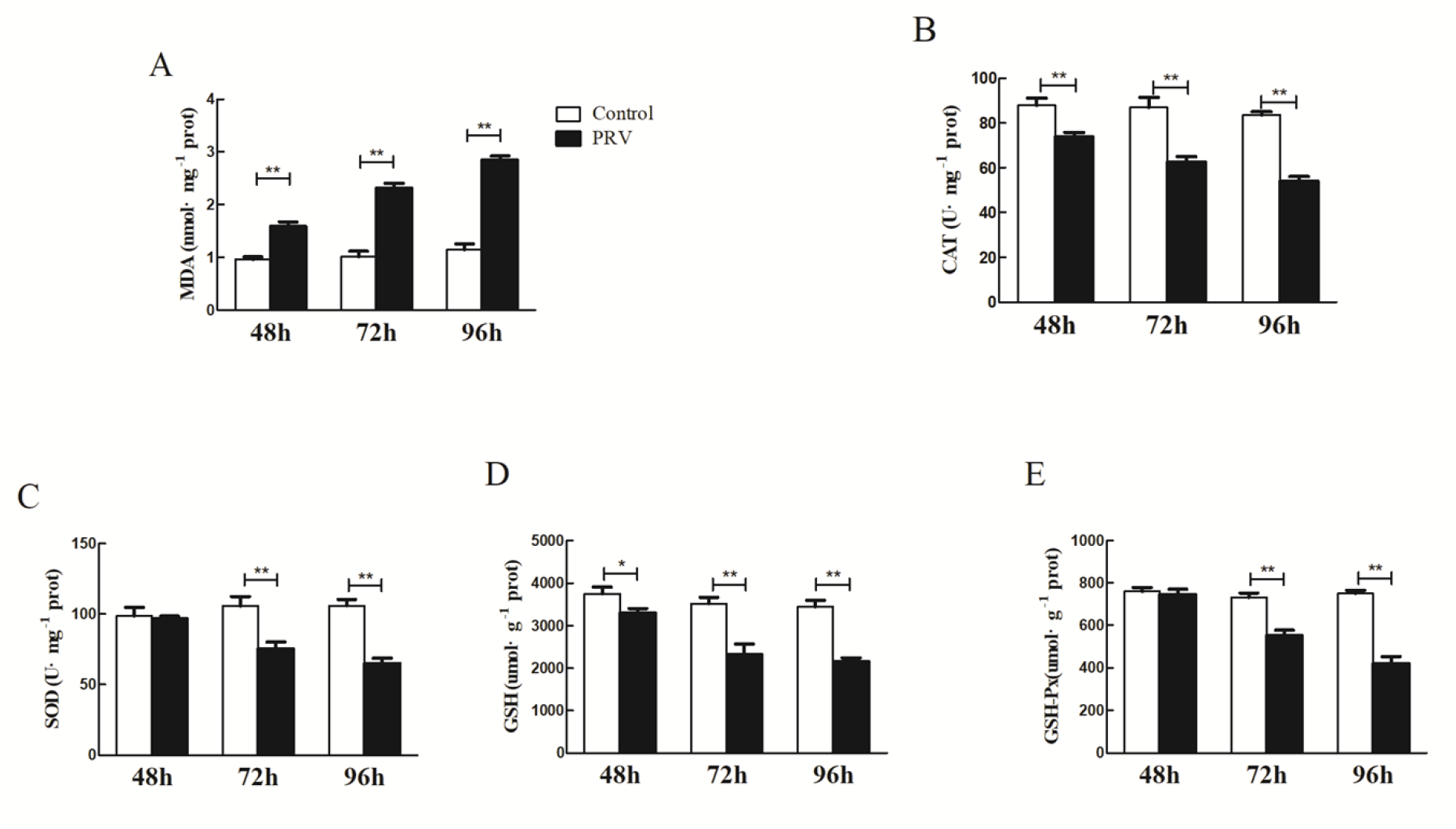
The effect of PRV infection mitochondrial oxidative stress-related factors (A)MDA; (B)CAT; (C) SOD; (D) GSH; (E) GSH-Px *\*p<0*.*05* versus mock infection, the same as below.

### 3.4 Detection of the splenocytes apoptosis

It can be seen from Figure 4, compared with their respective in the control, the proportion of early, lately and total apoptotic cells in the PRV infection group significantly increased in the infection period. In the 48 hpi time point, apoptotic cells were significantly increased (*p*<0 05).However, the apoptotic cells extremely increased at the 72 and 96 time points(*p*<0 01). PRV could promote the production of splenocytes apoptosis.

**Figure 4.**
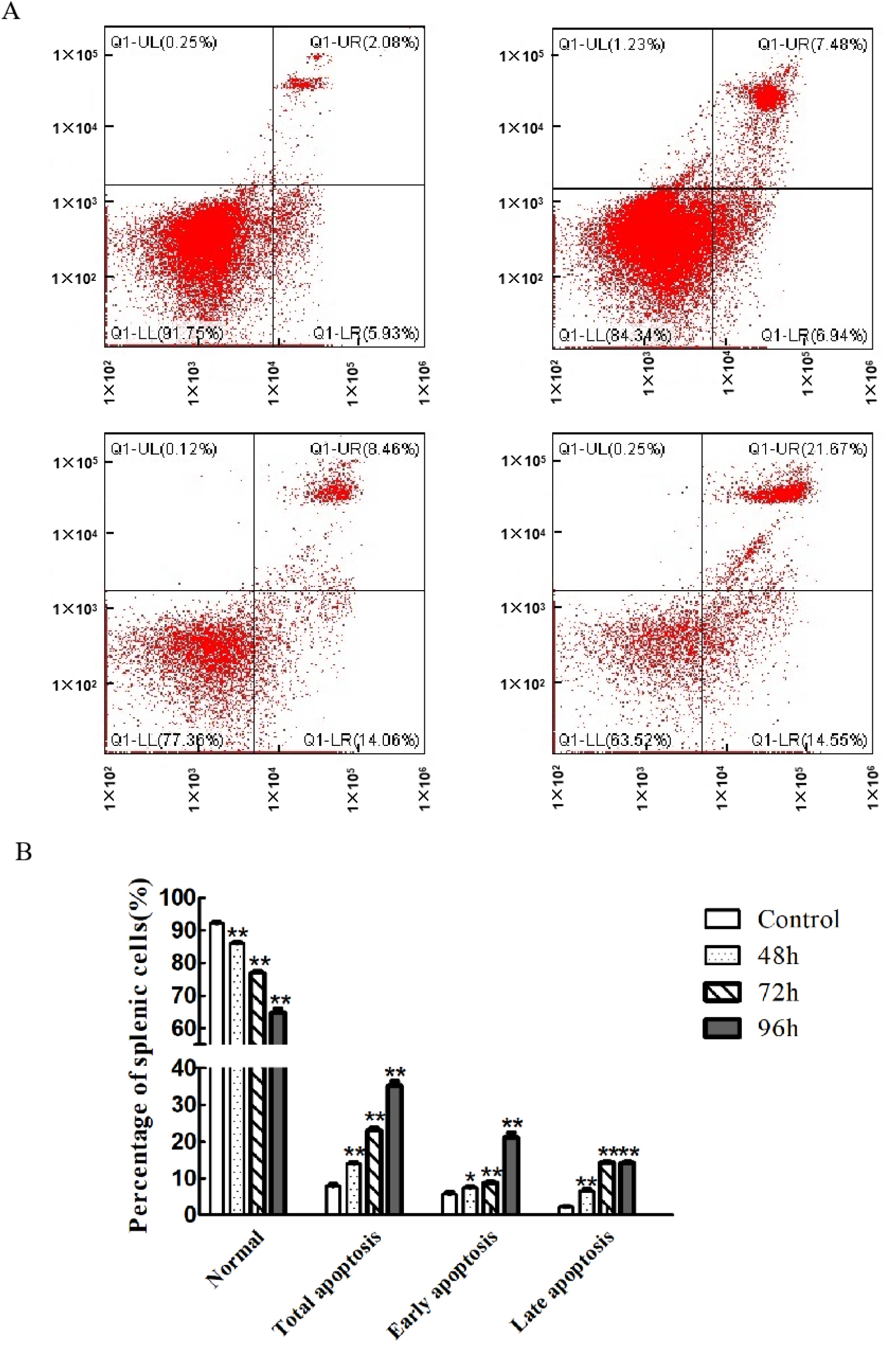
Effect of PRV on the percentage of splenocyte apoptosis (A) The results of apoptosis detected by flow cytometry; (B) ROS statistical analysis histogram

### 3.5 Relative mRNA expression levels related to apoptosis in the Spleen

From the results in Figure 5, Caspase-3 relative mRNA expression levels in the PRV infection group at 48 hpi, 72hpi and 96 hpi were all increased (*p*<0.05 or *p*<0.01) by comparing with their respective controls. Furthermore, qRT-PCR results demonstrated that Caspase-9 mRNA expression levels in the PRV infection group were also up-regulated at 72 hpi and 96 hpi groups (*p*<0.05 and *p*<0.01) compared to their respective control groups. Furthermore, the Bax expression levels of the PRV infection groups were not significant difference at the 48 hpi and 72 hpi time points compared to their respective controls(*p*>0.05). However, we observed a significant difference in Bax levels at the 96 hpi time point for PRV infection compared with the gene in the control group. In addition, the mRNA expression of Bcl-2 in the PRV infection group was statistically decreased at the 72 hpi and 96 hpi time points compared to their respective in the control(*p*<0.05 and *p*<0.01).

**Figure 5.**
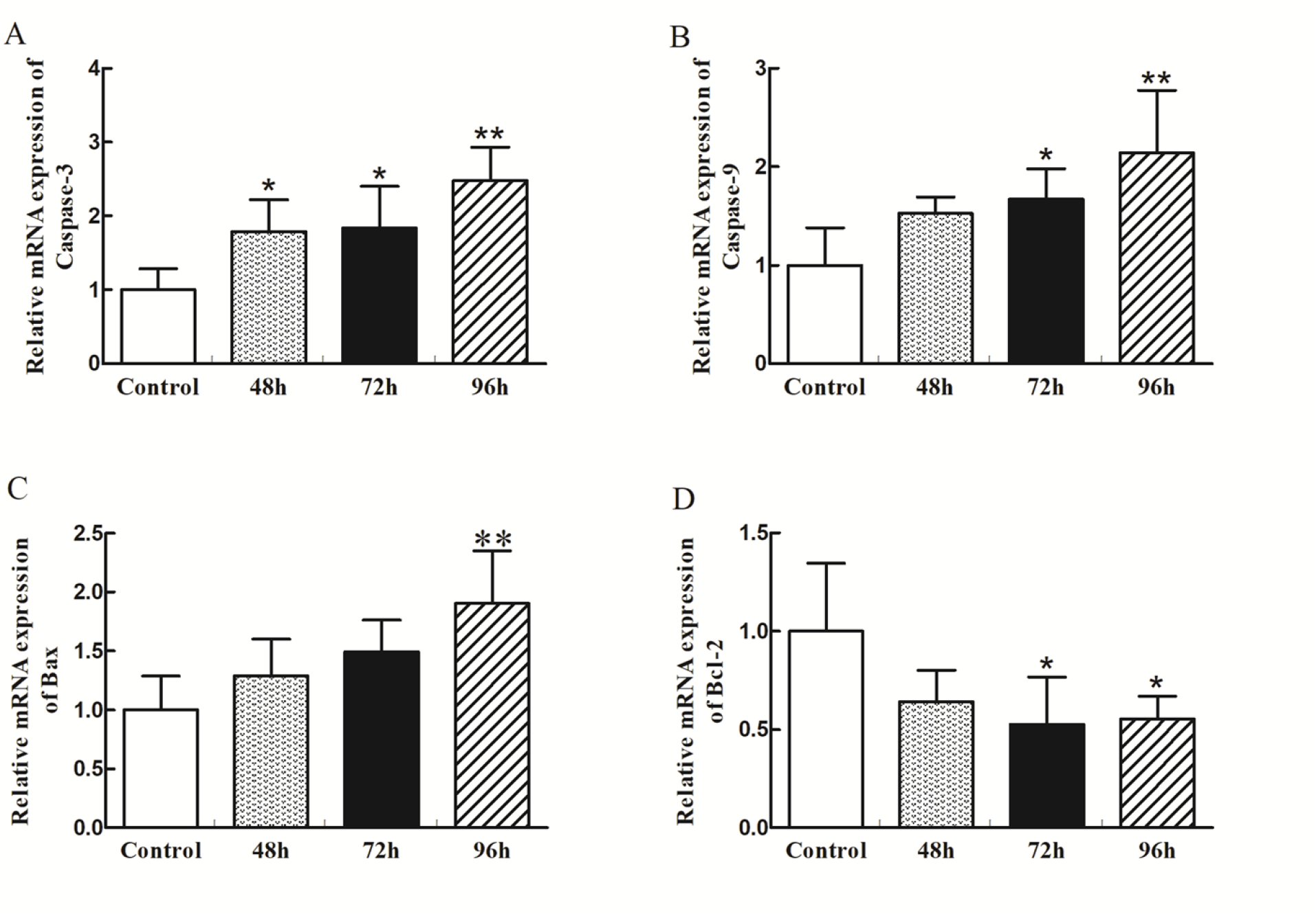
The relative mRNA expression of apoptosis-related factors caused by PRV

### 3.6 Effects of PRV on the proteins related to apoptosis in the Spleen

According to the results in Figure 6, Caspase-3 protein expression levels were significantly increased in the 48 hpi, 72hpi, and 96hpi PRV post-infection time points compared to their respective control group. In addition, compared with their respective control groups, the PRV infection group showed a significant up-regulation of Caspase-9 at 96 hpi post-infection time point. Moreover, the Bax expression in the PRV infection group remarkably rose at all the time points compared to their respective control groups. Furthermore, PRV down-regulated the protein expression of Bcl-2 in the 48 hpi, 72 hpi, and 96 hpi group by comparing them with the control groups.

**Figure 6.**
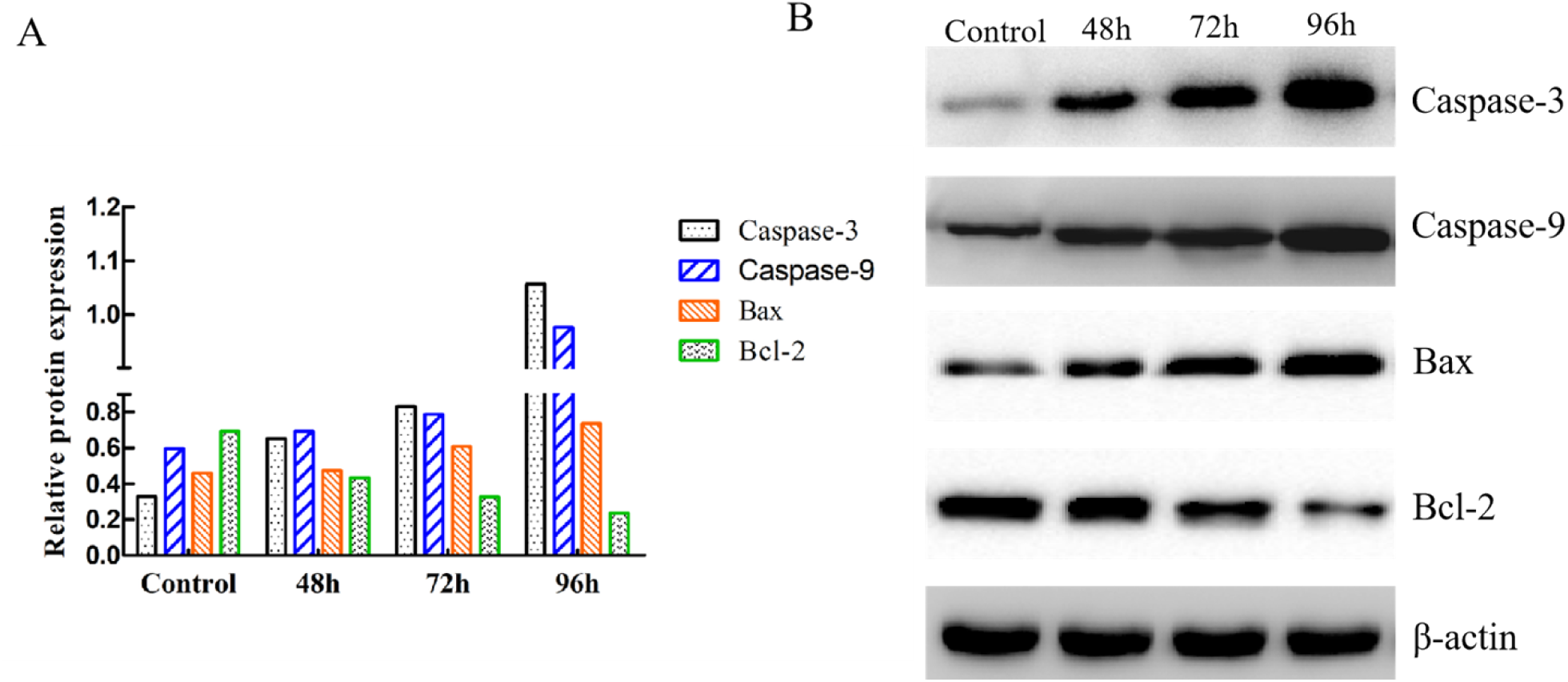
The expression levels of apoptosis-related factors induced by PRV in protein (A)The relative protein amount of Caspase-3, Caspase-9, Bax, and Bcl-2 to β-actin (B) Protein expression related to apoptosis detected by western blot

## 4. Discussion

The spleen is one of the major parenchymal organs affected by PRV. After PRV infection, the spleen mainly exhibits small white coagulative or lytic necrotic lesions[7]. In this work, In this experiment, we observed congestion and even inflammatory cell infiltration in the red pulp area of the spleen after PRV infection. Research has shown that PRV elevates the signaling pathways of p38-MAPK and JNK/SAPK, inducing tumor necrosis factor-α (TNF-α) mRNA levels and promoting of TNF-α secretion, which results in cell apoptosis[8]. An in vitro study showed that, a huge ROS was generated in the PK15 cells infected with PRV and this caused DNA damage and cell apoptosis[9]. The spleen is a substantial organ rich in mitochondria, an organelle that serves as the main source of ROS production. However, the mitochondria are also susceptible to ROS damage. ROS promotes the formation of lipid peroxidation, in which MDA is a key marker[10]. The oxidative stress induced by the out-of-balance between oxidants and antioxidants is an important feature of the occurrence of a series of viral infectious diseases[11]. It reported that HSV-1, a member of the PRV genus, causes GSH depletion and increases ROS and lipid peroxidation levels[12]. It has been reported in both cellular and animal models that HSV-1 causes host damage through oxidative stress[13]. In this experiment, after PRV was subcutaneously inoculated into mice, basing on the microscopic examination, we further observed that PRV promoted the production of ROS and MDA in the spleen of mice, reduce the production of non-enzymatic substances GSH, and at the same time down-regulated the expression of antioxidant enzymes i.e CAT, SOD, GSH-Px in the terms of gene levels, indicating that PRV can break the balance between splenocyte oxidation factors and antioxidant factors, hence induce oxidative stress in the spleen.

High levels of ROS are the main cause of cell apoptosis under physiological and pathological conditions. PRV induces the oxidative stress and damage of DNA, thus affecting the expression of pro-apoptotic signaling molecules and key regulatory molecules of cell cycle[9]. Apoptosis is a mechanism for maintaining balance in vivo, and has a pivotal role in viral infection. Several viruses directly induce host cell apoptosis at the stage of infection, which may be a pathological factor related to infection. It is known that PRV infection triggered suicide or apoptosis of infected cells[14]. Generally, apoptosis is activated mainly in the late stage of infection by PRV. Moreover, studies have shown that PRV triggers ROS production and causing of apoptosis in autophagic-damaged cells, which indicated that ROS may be participated in the linking of autophagy and apoptosis in cells infected by PRV[15]. Oxidative stress is associated with various cell death processes, including apoptosis, caused by viral infection. Hepatitis B virus (HBV) combines with the mitochondria through the C-terminal of X protein (HBX) to cause damage to mitochondrial DNA, and over-express the SIRT-1 gene for ROS production[16]. Interestingly, the Hepatitis C virus (HCV) causes damage to host cells in mitochondria by activating oxidative stress, and the damaged mitochondria are eliminated by way of autophagy[17]. In this work, our results indicated that the PRV-HLJ strain caused oxidative stress in the spleen of mice, thereby inducing splenocyte apoptosis. Hu Wei et.al[18] reported that the PRV Bartha strain up-regulated the expression of LMP2, LMP7, LMP10, and other immune-related protease sub-units. After infecting mice with the PRV-GXLB-2013 strain, they observed an increase in ROS production in the brain, spleen, and lung.[19]. From these results, we speculated that there is a certain connection between the immune response caused by PRV and an organ being in a state of oxidative stress, however, the specific mechanism is not yet clear, and requires further research.

Virus infection activates Caspase-3, leading to cell apoptosis[20]. Caspases, a series of enzymes in the cysteine protease family, plays a pivotal function in the maintaining of homeostasis and regulating of programmed cell death in vivo. Caspase-3 and caspase-9 have been widely classified based on their known role in cell apoptosis[21]. Caspase-3 activation is essential for the promotion of cell apoptosis. Bax and Bcl-2 are two regulators in the process of releasing cytochrome C and mitochondrial integrity. Research has shown that excessive levels of ROS have a crucial role in activating the mitochondrial-dependent apoptosis signal pathway, which promoted the release of cytochrome C, reduces levels of mitochondrial membrane potential (MMP), damages cell respiratory function, disturbances the imbalance of energy metabolism, and finally apoptosis[22]. Apoptosis is inevitably related to the change of MMP. The decline of mitochondrial membrane potential is an important landmark event for apoptosis[23].

The studies on the mechanism of PRV invasion are still unclear. The oxidative stress caused by the excessive generation of ROS in the spleen after being invaded by the virus is closely related to tissue damage. This was consistent with our experimental results which confirm that PRV infection caused spleen oxidative stress, thereby inducing splenocyte apoptosis. This study provides certain scientific data for further exploring the spleen damage caused by PRV and its pathogenic mechanism.

## 5. Conclusion

In conclusion, our study showed that Pseudorabies virus strain HLJ strain (MK080279.1) induces apoptosis in the spleen of mice via ROS production.

## Ethical Statement

Research on animal testing in this study was authorized by the Animal Ethics Committee of this College, all all experiments were performed in accordance with the approved guidelines and regulations.

## Authors’ Contributions

Wei Sun: Conceptualization and Writing-Original draft preparation; Shanshan Liu: done qRT-PCR and Western blot; Qingyan Wang, Yu Fan and Yi Yan: Data analysis and artwork making; Samuel Kumi Okyere: Supervision.

